# Performance and behavioral flexibility on a complex motor task depend on available sensory inputs in early blind and sighted short-tailed opossums

**DOI:** 10.1101/2020.05.12.091108

**Authors:** Mackenzie Englund, Samaan Faridjoo, Chris Iyer, Leah Krubitzer

**Affiliations:** Department of Psychology, University of California, Davis. 135 Young Hall, 1 Shields Ave., Davis, CA 95616, USA; Department of Molecular and Cellular Biology, University of California, Davis. 149 Briggs Hall, 1 Shields Ave., Davis, CA 95616, USA; Symbolic Systems Program, Stanford University, 460 Margaret Jacks Hall, 450 Serra Mall, Stanford, Ca 94305, USA; Center for Neuroscience, University of California, Davis. 1544 Newton Ct., Davis, CA 95618, USA

## Abstract

The early loss of vision results in a reorganized visual cortex that processes tactile and auditory inputs. Recent studies in the short-tailed opossum (*Monodelphis domestica)* found that the connections and response properties of neurons in somatosensory cortex of early blind animals are also altered. While research in humans and other mammals shows that early vision loss leads to heightened abilities on discrimination tasks involving the spared senses, if and how this superior discrimination leads to adaptive sensorimotor behavior has yet to be determined. Moreover, little is known about the extent to which blind animals rely on the spared senses. Here, we tested early blind opossums on a sensorimotor task involving somatosensation and found that they had increased limb placement accuracy. However, increased reliance on tactile inputs in early blind animals resulted in greater deficits in limb placement and behavioral flexibility when the whiskers were trimmed.

## INTRODUCTION

Navigation through complex space requires a tight coupling between incoming sensory input and motor output (Abbruzzese & Berardelli, 2003; Ferezou et al., 2007). The ability of an animal to make precise, sensory-guided movements while actively exploring the environment depends on the sensory information available, the structure of the peripheral epithelia, the morphology of the body, and the neural circuits involved in integrating sensory information from different modalities. Because evolution has produced species who rely on some senses over others, it is important to look at animals that have different body morphologies and rely on different combinations of sensory input to appreciate general principles of nervous system structure and function that allow mammals to adapt movement strategies in a constantly changing environment. For example, cats and humans use vision to guide complex locomotion (Drew & Marigold, 2015; McVea et al., 2009); bats rely on echolocation to navigate three-dimensional space (Moss & Surlykke, 2010); and rodents employ tactile inputs relayed through the whiskers to navigate terrestrial and arboreal habitats (Grant et al., 2009; Mitchinson et al., 2011). While different species often rely heavily on one sense, there is accumulating evidence that the developing neocortex is not as strictly constrained by its evolutionary history as previously thought, since the loss of sensory input early in development leads to a massive restructuring of the neocortex based on the remaining sensory inputs (Bell et al., 2019; Cecchetti et al., 2016; Kupers & Ptito, 2014; Renier et al., 2014).

Studies of early vision loss in animal models and humans demonstrate that the neocortex is capable of remarkable functional and anatomical plasticity, and this plasticity appears to support important sensory-mediated behaviors. For example, if visual input is lost in developing mice, rats, and hamsters, neurons in visual cortex (V1) instead respond to deflections of the whiskers and to auditory stimuli such as tones and clicks (Izraeli et al., 2002; Piché et al., 2007; J. Toldi et al., 1990). In addition, cortical and subcortical connections of V1 undergo significant alterations that are shaped by the remaining sensory inputs. In rats, early vision loss results in a decrease in connectivity between visual thalamic relay nuclei and visual cortex (LGN-V1), and an increase in connectivity between association thalamic nuclei and visual cortex (LP-V1) (Négyessy et al., 2000). Similar alterations to thalamic relay and association nuclei have been observed in cats enucleated at birth (Berman, 1991). Cortico-cortical connections are also altered by the early loss of vision. Rats enucleated at birth show increased variability and divergence of intra- and interhemispheric connections between V1 and extrastriate cortex (Bock & Olavarria, 2011; Laing et al., 2012). In mice enucleated at birth, lipophilic dye tracing experiments show that the early loss of vision alters cortico-cortical connections prior to would-be eye opening, such that projections from S1 to V1 are more broadly distributed in blind mice compared to sighted controls (Dye et al., 2012; Kozanian et al., 2015).

Work from our laboratory in short-tailed opossums (*Monodelphis domestica*) has demonstrated that removing all visual input prior to the onset of spontaneous activity from the retina results in massive cross-modal changes in the functional organization and connections of what would have been visual cortex (Kahn & Krubitzer, 2002; Karlen & Krubitzer, 2009). Similar to rodents, neurons in the reorganized visual cortex of early blind opossums respond instead to deflections of the whiskers and broadband auditory clicks (Karlen et al., 2006). The spared sensory systems are also affected, as early blind opossums exhibit changes in the cortical connections of S1, as well as changes in receptive field size and response properties of neurons in the whisker representation of S1 (Dooley & Krubitzer, 2019; Ramamurthy & Krubitzer, 2018).

Yet only a few studies have measured the behavioral manifestations of cross-modal anatomical and functional plasticity in animal models following early loss of vision. For example, cats deprived of visual input through eyelid suture show enhanced sound localization abilities (Rauschecker, 1995; Renier et al., 2014), and preliminary data from our own laboratory shows that enucleated opossums have better texture discrimination (Dodson et al., 2017).

Studies in humans also demonstrate behavioral, anatomical and functional cross-modal effects following early loss of vision. For example, primary visual cortex is recruited during tactile and auditory discrimination tasks, including braille reading and pitch detection (Collignon et al., 2011; Gizewski et al., 2003; for review Ricciardi et al., 2014; Van der Lubbe et al., 2010). This is accompanied by better texture and pitch discrimination, and shorter reaction times to detect auditory and somatosensory spatial targets (for review Bell et al., 2019; Goldreich & Kanics, 2003; Norman & Bartholomew, 2011; Postma et al., 2016). Similar to studies in animals, diffusion tensor imaging (DTI) studies in congenitally blind humans show a reduction in the size and connectivity of V1, and an increase in the connectivity between S1 and V1 (Li et al., 2013; Ptito et al., 2008; Shu et al., 2009). The stronger connections between the spared sensory cortical areas and V1 seems to be restricted to congenitally blind individuals; those with late onset blindness or partial vision show weaker connectivity between S1 and V1 (Burton et al., 2014). Interestingly, regardless of the age of onset, resting state fMRI and TMS studies have shown that there is an increase in connectivity between V1 and regions of the brain associated with memory and spatial processing (e.g. hippocampus, posterior parietal cortex) in blind compared to sighted individuals (Burton et al., 2014; Liu et al., 2007; Pelland et al., 2017; Rimmele et al., 2019; Wang et al., 2014). Given the overwhelming evidence in both animal models and humans that early vision loss results in drastic cross-modal changes to sensory and association cortical areas, it is surprising that few studies have investigated the behavioral outcomes that result from these neural changes, and none have quantified how kinematics, strategy, and performance during complex navigation are altered in tandem.

In fact, we know very little about the effects of early sensory loss on some of the fundamental behaviors of mammals: locomotion and navigation. One study in rats enucleated at birth found that blind animals perform a maze-running task more quickly, and have alterations in the receptive-field size of neurons in barrel cortex (Toldi et al., 1994a), while a finger-maze study in blind humans found no difference in performance compared to sighted participants (Gagnon et al., 2012). Other studies have found that blind individuals can create mental maps of space more efficiently than sighted controls (for review: Schinazi et al., 2016). However, in many spatial navigation tasks, blind individuals are at a disadvantage. This is due to the ability of sighted subjects to integrate space over large distances using vision, while blind participants are restricted to acquiring one meter of spatial information per cane movement (Schinazi et al., 2016). Furthermore, many spatial tasks constrain both blind and sighted participants to adopt similar strategies (e.g. by forcing sighted participants to wear a blindfold), which predisposes individuals of a given condition to be at a selective disadvantage (Gagnon et al., 2012; Haber et al., 1993). In a recent review, Shinazi and colleagues raise this as an important concern for understanding the differing abilities of blind and sighted individuals, suggesting that performance and strategy be studied in tandem (Schinazi et al., 2016).

The goal of the present investigation was to assess differences in both performance and strategy between early blind and sighted opossums on a naturalistic sensorimotor task. We also sought to determine the extent to which animals rely on the remaining available sensory information by removing sensory driven input from the whiskers. To accomplish this, we utilized recently developed, marker-less tracking algorithms to analyze performance, locomotor patterns, and movement strategies captured on video, in short-tailed opossums bilaterally enucleated at postnatal day 4; prior to the onset of spontaneous retinal activity and the formation of thalamic and cortical visual pathways. As *Monodelphis* is among mammals that use the whiskers for active sensing (known as whisking), we used a classic sensorimotor task known to involve somatosensory detection/discrimination: the variable ladder-rung walking task (Metz & Whishaw, 2009; Mitchinson et al., 2011). This task is sensitive to minute differences in motor control, and has been used to assess motor deficits in peripheral and neurologic disease models (Antonow-Schlorke et al., 2013; Metz & Whishaw, 2002; Schönfeld et al., 2017). Our results link previously observed cross-modal changes in the connections and functional organization of the neocortex to changes in sensorimotor behavior, and the kinematics associated with complex navigation.

## RESULTS

In the following results we compared both performance and movement strategy in early blind (EB) and sighted control (SC) animals while performing the ladder rung task. Data is described as Mean ± SEM. Statistics are reported as Adjusted R-Squared (R^2^adj), and the F-Statistic is presented with degrees of freedom of the model, F(N). Delta (Δ) indicates the difference between pre whisker-trim and post whisker-trim means within early blind (ΔEB) and sighted opossums (ΔSC).

### Early blind opossums outperform sighted controls in variable ladder rung walking, but show similarities in crossing strategy

To determine differences in ladder rung walking performance between early blind and sighted opossums, foot faults on nine variable ladder rung patterns were scored by two independent observers. On average, early blind opossums (EB) had a per-run error rate of 4.74 ± 0.88% in the light condition, and an error rate of 4.59 ± 0.62% in the dark condition. Sighted controls (SC) had an error rate of 10.28 ± 1.13% in the light condition, and 8.61 ± 1.01% in the dark condition. Using a linear model testing for interactions between experimental group and lighting condition, we found a significant main effect for sightedness on total error, with early blind opossums committing fewer errors than sighted controls (R^2^adj = 0.32, F(8) = 6.62, p = 0.007) (Figure 2A). Under the same model, we found no significant effect of lighting (p =0.718), with animals in both experimental groups performing similarly in the light and dark (Supplemental Figure 1C). Further, we found no effect of sex on performance (p = 0.086). We found no differences in performance across the patterns of rung placement, nor in the order in which rung patterns were completed (light or dark first) (R^2^adj =0.09, F(3) = 8.00, p = 0.367, p = 0.207) (Supplemental Figure 1A). As there was no significant effect for lighting, a reduced model was used to test the individual contributions of forelimb and hindlimb error to total error rate.

**Figure 1.**
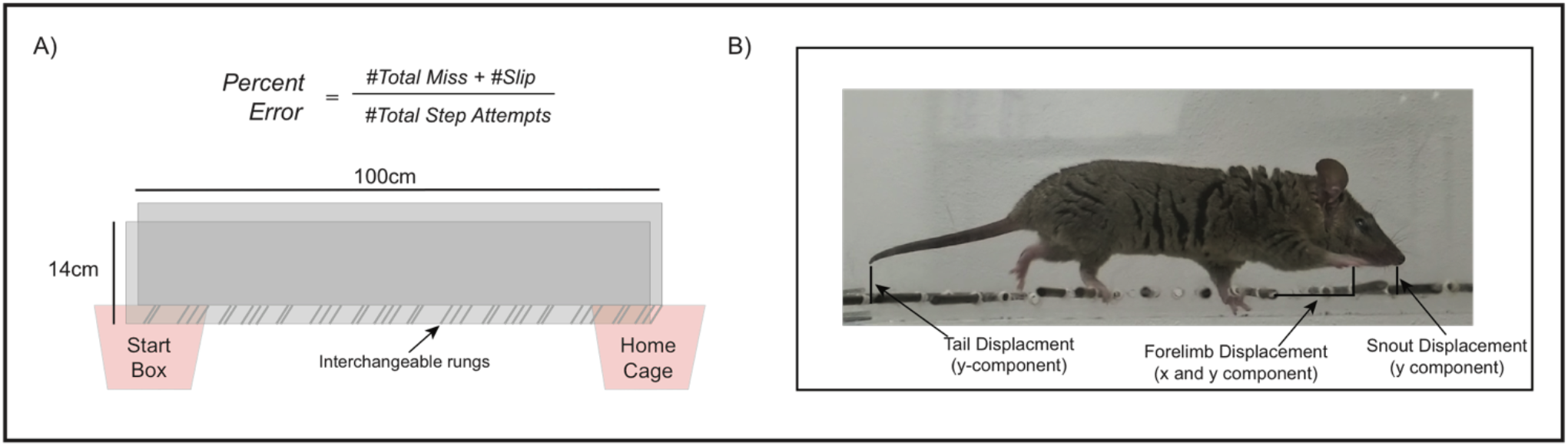
Variable Ladder Rung Walking. (A) Equation for error rate (top) and ladder rung apparatus (bottom). During a single trial, opossums move from a neutral start box (left) to their home cage (right). (B) Image of a sighted opossum on the ladder rung apparatus, illustrating aspects of posture chosen for analysis.

**Figure 2.**
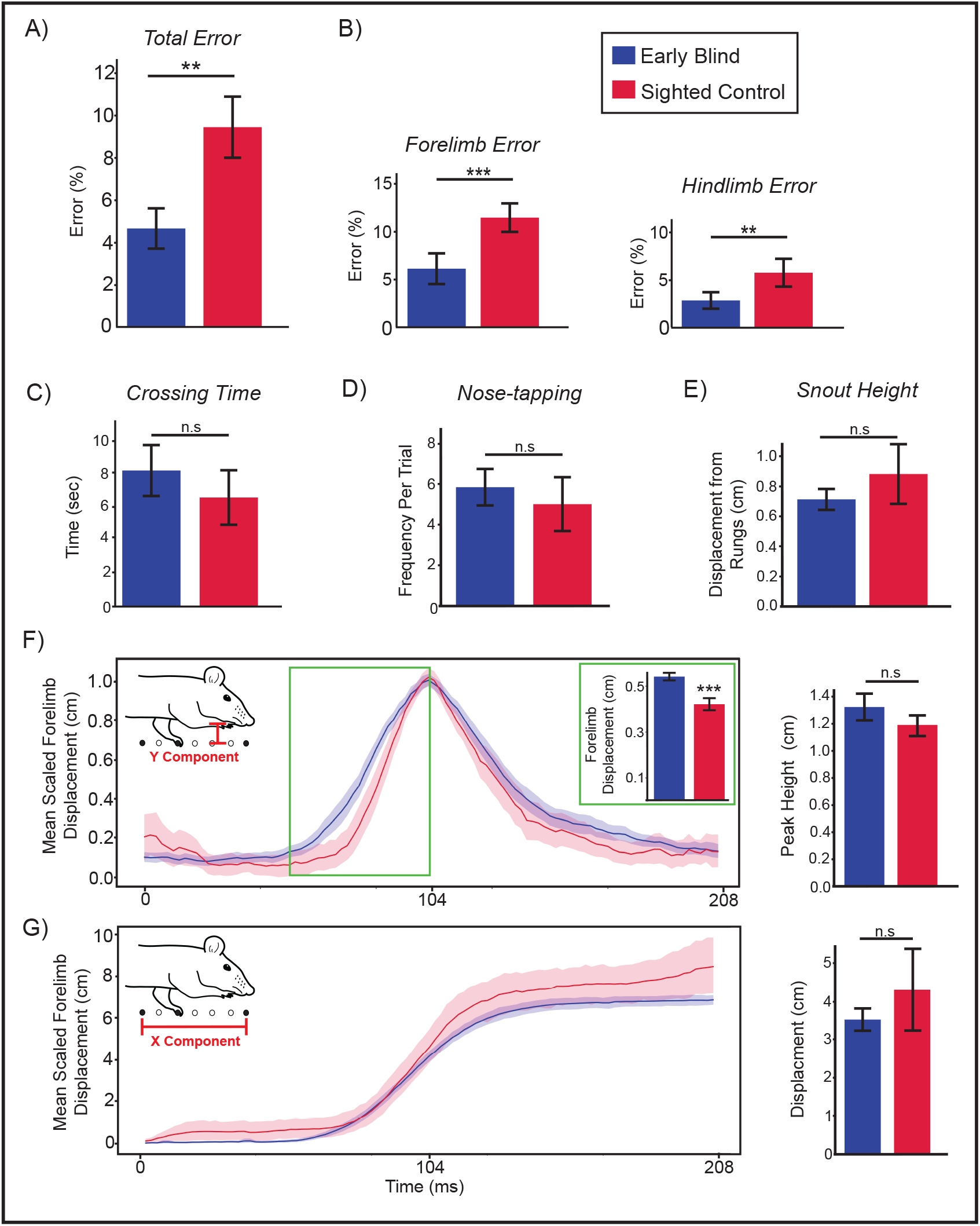
Early blind opossums outperform sighted controls in variable ladder rung walking, but show similarities in crossing strategy. (A) Bar graphs showing that early blind opossums commit significantly less error than sighted opossums (p = 0.007). (B) Bar graphs of average forelimb and hindlimb error. On average, early blind animals commit significantly less forelimb (p = 0.004) and hindlimb (p = 0.008) error than sighted controls. (C) We found no significant difference in crossing time (p = 0.27), (D) average number of nose-taps per trial (p = 0.626), or (E) snout height during correct forelimb placements between conditions (p = 0.806). (F) Line graphs depicting the average y-component (step height) of forelimb trajectories during correct placements (n = 99 EB; 56 SC) of 8 animals (n = 4 EB; n = 4 SC), scaled by average peak height per animal (inset image, left panel). The green square and inset bar graph denotes the region of non-overlap of 95% confidence intervals, where early blind animals are quicker to lift their right forelimb off of a rung (p < 0.001). Bar graph shows quantification of average y-component forelimb trajectory as average peak height (right). Note: bar graph does not include scaling to illustrate there is no statistical difference in peak height between conditions prior to scaling (p = 0.267). (G) Line graphs depicting forelimb displacement in the x-direction (stride length). Quantification of average displacement shows no difference in the average x-component of forelimb movement between early blind and sighted opossums (p = 0.405) (right). All bar graphs are presented as between-group averages. Error bars are presented as 95% bootstrapped confidence intervals.

Early blind opossums committed significantly less forelimb errors (6.13 ± 0.87%) than sighted animals (11.53 ± 0.87%), (R^2^adj = 0.32, F(4) = 11.81, p = 0.004) (Figure 2B). Additionally, early blind opossums committed significantly less hindlimb errors (2.88 ± 0.47%) than sighted animals (5.79 ± 0.76%), (R^2^adj = 0.10, F(4) = 3.71, p = 0.008) (Figure 2B). Thus, in the absence of vision, early blind opossums outperformed their sighted counterparts, making significantly fewer errors during rung walking whether fore- and hindlimbs were considered separately or together.

To investigate the crossing strategies and kinematics that may underlie differences in performance, we quantified crossing time and nose tapping behavior, and used DeepLabCut marker-less tracking in a randomly-chosen subset of animals (n = 8) to track the position of the snout, limbs, and tail, as animals crossed the ladder. We found no significant effect of crossing time on performance (R^2^adj = 0.35, F(4) = 10.71, p = 0.23), and no significant difference in crossing time between early blind (7.85 ± 0.72s) and sighted animals (6.64 ± 0.9s) (R^2^adj = 0.24, F(4) = 7.67, p = 0.27) (Figure 2C). We also found no significant difference in the average frequency of nose taps per-trial between blind and sighted animals (R^2^adj = 0.34, F(4) = 7.57, p = 0.63) (Figure 2D), or the average height at which animals held their snout during correct placements (R^2^adj = 0.60, F(4) = 6.46, p = 0.84) (Figure 2E). These results show that animals in both groups take similar amounts of time to cross the ladder and implement some of the same strategies while doing so, but early blind animals show increased performance.

Next, we quantified average forelimb trajectories during correct placements in both groups (n = 155 forelimb motions). Average peak height did not differ between groups (R^2^adj = 0.24, F(4) = 7.67, p = 0.27) (Figure 2F). However, during forelimb motion, early blind animals are relatively quicker to lift their forelimb compared to sighted control animals (R^2^adj = 0.72, F(4) = 1646.0, p < 0.001) (Figure 2F; Green Rectangle). This was quantified as the average height of the limb during the second quartile of a forelimb motion (52 – 104ms). Surprisingly, average forelimb peak height significantly predicted average forelimb error, with higher peaks indicating better performance (R^2^adj = 0.78, F(4) = 13.06, p = 0.028). While one might expect larger movements to be correlated with more errors, peak height may be an indicator of an animal’s confidence in its estimation of where the next rung is located. In the X-direction (step length), both early blind and sighted opossums exhibit similar forelimb kinematics (i.e. similar step lengths), possibly constrained by the finite spacing of the rungs (Figure 2G). Thus, apart from differences in when the forelimb was lifted off of a rung, there was no overall difference in the behavioral strategy shown by the metrics that we used. However, it should be noted that differences in the kinematics of other body parts that we did not quantify, particularly in how the position of the whiskers relates to the position of the forelimbs, may also contribute to differences in performance. The data presented below on whisker trimming indicate that this may indeed be the case.

### Whisker trimming causes deficits in performance in forelimb but not hindlimb placement. Performance deficits and limb trajectories are altered to greater extents in early blind animals

Work from our laboratory has recently shown that neurons in the whisker representation of S1 have smaller receptive fields and increased selectivity (i.e. greater discriminability) (Ramamurthy & Krubitzer, 2018). Thus, we sought to examine the role of the whiskers during variable ladder rung walking. To accomplish this, we trimmed the mystacial and genal vibrissae, and re-tested animals on variable patterns over the course of two additional testing days. We predicted that both early blind and sighted animals would exhibit significant performance deficits, as short-tailed opossums are among animals who actively whisk (Grant et al., 2013; Mitchinson et al., 2011). Accordingly, we found a significant main effect for trimming of whiskers on overall performance in both early blind and sighted animals (R^2^adj = 0.31, F(4) = 11.62, p < 0.001, ΔEB = +5.57% error, ΔSC = +2.90% error) (Figure 3A). Moreover, increases in error due to whisker trimming are driven by increases in forelimb error (R^2^adj = 0.32, F(4) = 11.81, p < 0.001, ΔEB = +8.81% error, ΔSC = +5.92% error) but not hindlimb error (R^2^adj = 0.10, F(4) = 3.71, p = 0.40, ΔEB = +1.09% error, ΔSC = +0.65% error) (Figure 3B). Whisker trimming caused a two-fold increase in total error in early blind opossums, increasing their error rate to similar levels to those observed in sighted animals with whiskers, highlighting their heavy reliance on this spared sensory system. Whisker trimming also significantly increased error in sighted animals, but only by a 1.3-fold change. We found that overall, average forelimb error significantly predicted average hindlimb error (R^2^adj = 0.32, p < 0.001 for animals with whiskers, and R^2^adj = 0.10, p = 0.03 for whisker trimmed animals) (Figure 3C), illustrating the relationship between forelimb and hindlimb movement in precision quadrupedal locomotion, as animals that were accurate forelimb placers were also accurate hindlimb placers.

**Figure 3.**
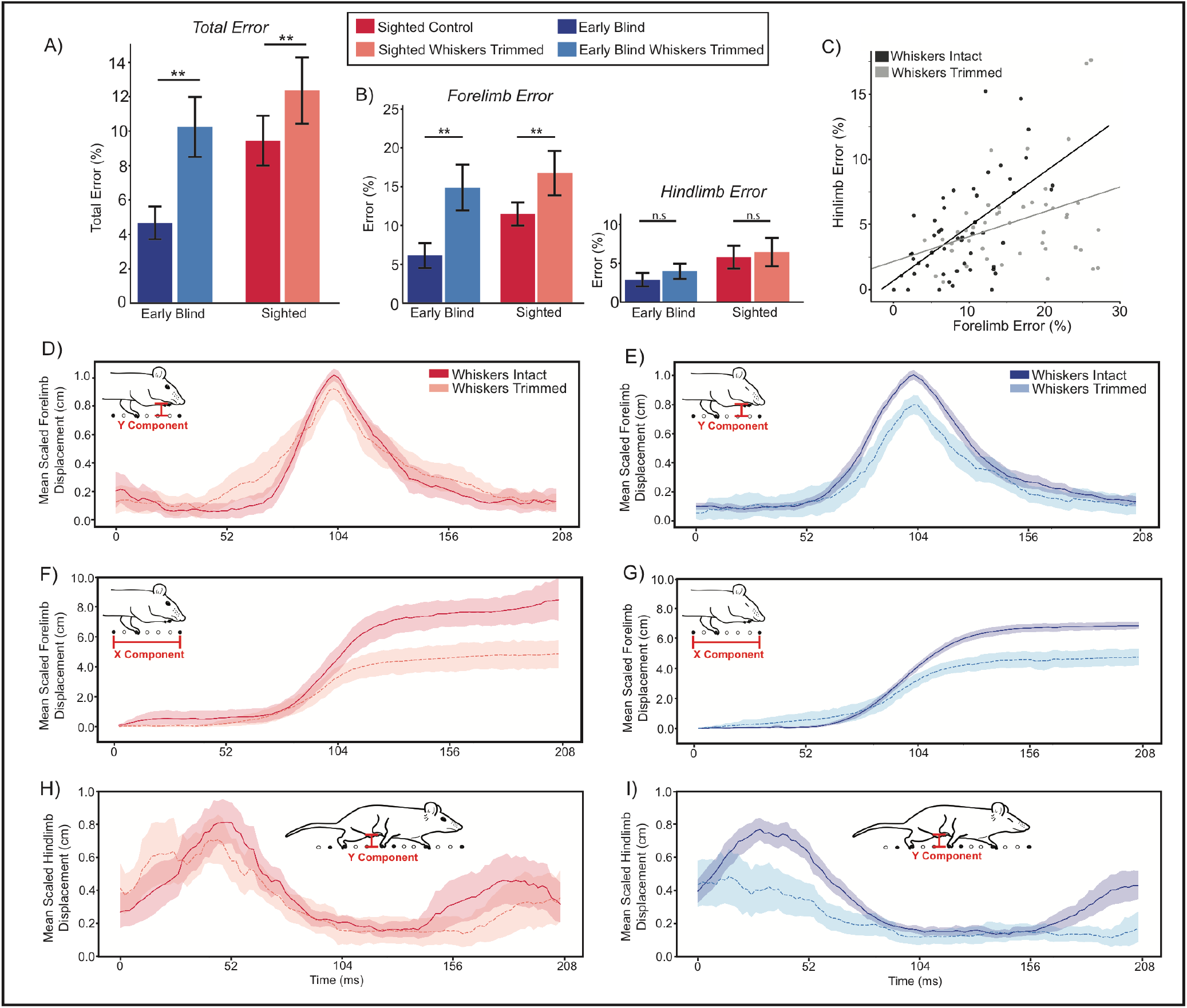
Whisker trimming causes deficits in performance in forelimb but not hindlimb placement. Performance deficits and limb trajectories are altered to greater extents in early blind animals. (A) Bar graph showing average total error before and after whisker trimming in early blind (blue) and sighted (red) opossums. Whisker-trimmed averages are represented by lighter shades (early blind whisker-trimmed: light blue; sighted whisker-trimmed: light red). A main effect was found for the presence of whiskers on performance (p < 0.004), with whisker trimming leading to increases in error. (B) Forelimb (left) and hindlimb (right) error before and after whisker trimming. Collapsing across lighting condition, the reduced model shows a significant main effect for increases in forelimb (p < 0.001) but not hindlimb (p < 0.397) error due to whisker trimming. (C) Scatter plot with fit lines showing that by animal, average forelimb error significantly predicts average hindlimb error in the presence (p < 0.001) or absence (p < 0.03) of whiskers. (D, E) Line graphs depict average y-component forelimb trajectories before (solid lines) and after whisker trimming (dashed lines) in sighted (red) and early blind opossums (blue). Whisker trimming reduces step height to a significant extent (p < .019). (F, G) Line graphs depicting forelimb displacement in the x-direction (stride) before and after whisker trimming. Whisker trimming reduces step width in both conditions. (H, I) Line graphs depict average y-component hindlimb trajectories before and after whisker trimming. Whisker trimming results in shallow hindlimb movements for blind and sighted animals. Overall, trajectories of the limbs are altered to a greater extent in early blind opossums. All bar graphs are presented as averages. Error is presented as bootstrapped 95% confidence intervals.

Next, to assess how the absence of tactile sensory information provided by the whiskers altered limb placement, we compared forelimb and hindlimb trajectories during correct forelimb placements before and after whisker-trimming using data derived from DeepLabCut marker-less tracking. We found that whisker trimming differentially altered the trajectories of forelimb motion in early blind and sighted animals, such that sighted animals exhibited little reduction in peak height while whisker-trimmed early blind animals show a significant reduction in forelimb peak height during correct placement (R^2^adj = 0.40, F(4) = 3.46, p = 0.02, ΔEB = −0.27 cm, ΔSC = −0.1 cm) (Figure 3D; Figure 3E; Supplemental Figure 1D). The 2.5-fold differential reduction in average forelimb peak height between groups demonstrates the greater effect of whisker trimming on early blind opossums’ sensorimotor coordination. The X-component (step length) of forelimb trajectories during correct placements was also altered due to whisker trimming, resulting in shorter steps for both groups (ΔEB = −0.8 cm, ΔSC = −1.69 cm) (Figure 3F; Figure 3G). Likewise, hindlimb trajectories in both groups became shallower after whisker trimming (Figure 3H; Figure 3I). Interestingly, the average reduction in hindlimb peak height matched the reduction in forelimb peak height in EB opossums exactly (ΔEB = −0.27 cm). Again, a less extreme reduction in hindlimb peak height was observed in sighted controls after whisker trimming (ΔSC = −0.23 cm) (Supplemental Figure 1D). This data highlights the importance of whiskers for forelimb placement, and further, the importance of forelimb placement in guiding hindlimb placement. In sum, removing a heavily relied upon sense, especially in early blind animals, results in changes to performance and locomotor trajectories.

### Whisker trimming alters forelimb dynamics by causing variable motions and different movement types

As we found that whisker trimming resulted in changes to forelimb trajectories, we sought to determine the underlying sources of these changes. Observation from video and trajectory plots indicates that variance in forelimb movements may increase after whisker trimming. Therefore, we assessed the variance of the Y-Component (step height) of forelimb trajectories between early blind and sighted opossums, with and without whiskers. As expected, the lowest variance in all groups was at the beginning, peak, and end of a forelimb movement (Figure 4A). While this is most likely due to the fact that peaks were aligned for analysis and that ladder rungs were at fixed intervals, restrictions on the musculoskeletal system and muscle synergies also contribute to highly stereotyped movements during locomotion. Variance of forelimb movements was not correlated with the number of strikes analyzed (R^2^adj = 0.01, F(1) = 1.12, p = 0.31). However, by animal, higher variance of correct forelimb placements was found to significantly predict average hindlimb error, again indicating that animals may be less certain of an upcoming rung’s position (R^2^adj = 0.30, F(1) = 7.47, p = 0.02).

**Figure 4.**
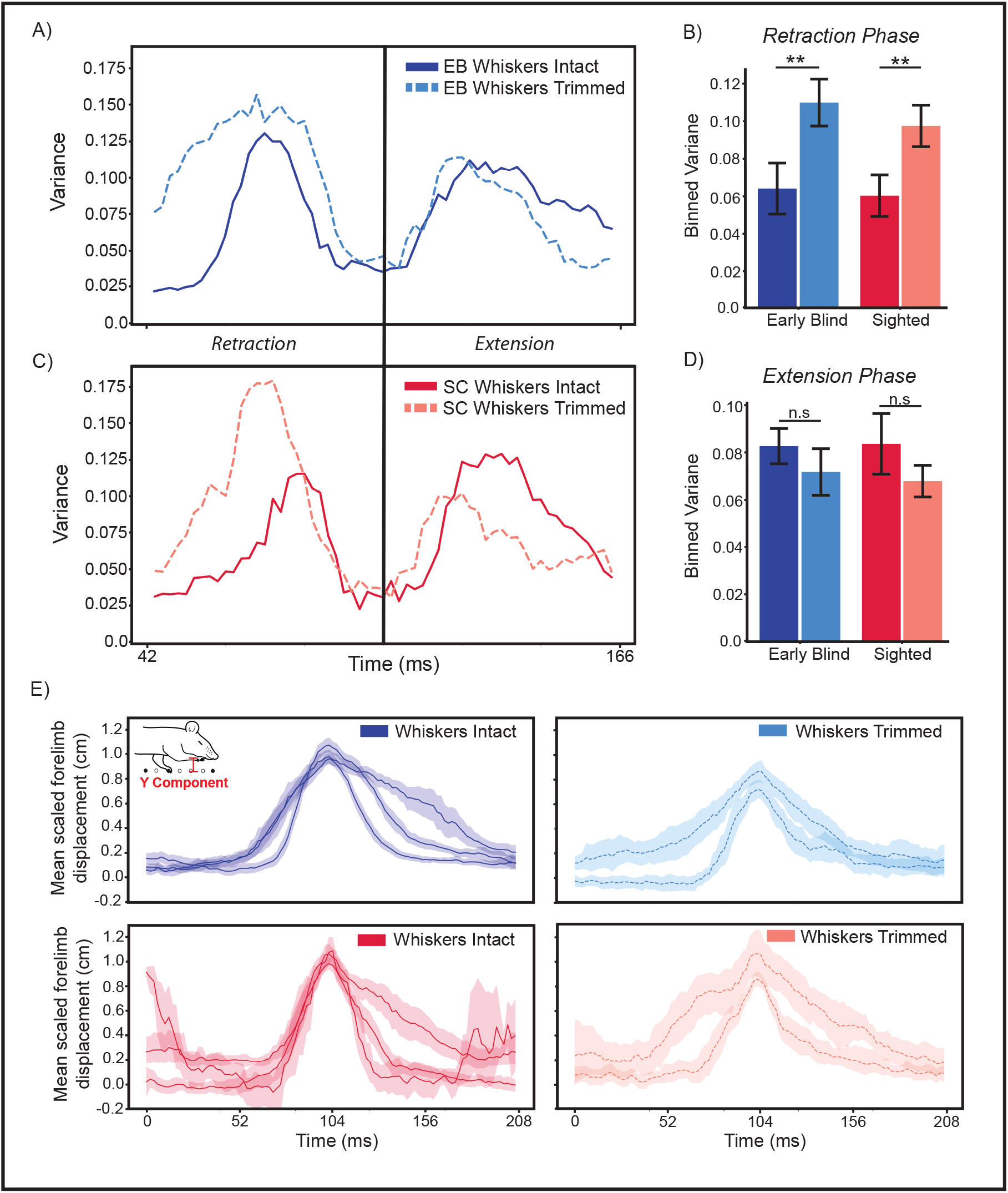
Whisker trimming results in more variable forelimb trajectories and different types of stereo-typical movements in both early blind and sighted animals. (A, C) Line graphs of the average variance of y-component forelimb trajectories across all motions (n = 155) before (solid lines) and after whisker trimming (dashed lines) in early blind (blue) and sighted (red) opossums. The vertical black line denotes the separation between retraction and extension phases. (B, D) Bar graphs show binned variance during retraction (B) and extension (D) phases. Binned variance is the average variance of the entire retraction or extension phase. Whisker trimming significantly increases variance during the retraction (p < 0.001), but not extension (p = 0.21) phase of forelimb movement. (E) Stereotyped movements of early blind and sighted opossums before and after whisker trimming provided by K-means clustering. The elbow point and silhouette coefficient were used to determine the number of clusters (see Supplementary Materials). Stereotypical movements are similar in early blind (top left) and sighted (bottom left) opossums. Whisker trimming alters stereotypical movements in similar ways in early blind (top right) and sighted (bottom right) opossums.

We then binned our analyses into a pre-peak retraction (frames 20 - 50), and a post peak extension (frames 50 - 80) phase. We excluded the first and last 20 frames of motion from this analysis to ensure frames where the forelimb was potentially resting on a rung were excluded. The retraction phase is noted to be the swing portion of the forelimb movement (Jacobs et al., 2014). This part of the movement begins on a ladder rung. The elbow then moves from full extension to full flexion at the peak of the motion. The extension phase is noted to be the swing-to-stance portion of the limb movement, where the elbow goes from flexion to extension, ending with the forepaw being placed on an upcoming rung. While there was no difference in the overall variance between early blind (0.063 ± 0.007cm) and sighted (0.060 ± 0.005cm) opossums (R^2^adj = 0.22, F(3) = 11.51, p = 0.23), whisker trimming significantly increased variance in both groups during the retraction phase of a forelimb movement (R^2^adj = 0.22, F(3) = 11.51, p < 0.001, ΔEB = +0.046, ΔSC = +0.036) (Figure 4B).

Conversely, whisker trimming did not significantly affect variance in the extension (placement) phase of forelimb movement (R^2^adj = 0.042, F(3) = 2.69, p = 0.21, ΔEB = −0.011, ΔSC = −0.015) (Figure 4A – Figure 4D). Together, this data shows that whiskers are critical for quick and accurate detection of future forelimb placement locations, but that the guidance of the motion is not affected once the opossum has already detected a rung.

Next, to characterize stereotypical movement types between groups, we used K-Means clustering to classify forelimb step height waveforms into stereotypical movements. *A priori*, we sorted waveforms by sightedness and whisker presence to detect differences within groups, and used the elbow point method and highest silhouette coefficient (SCoef) to determine the number of clusters used for each experimental condition. For sighted animals, both methods converged on three clusters (SCoef_3_ = 0.24) (Figure 4E; Supplementary Figure 2). For early blind animals, the elbow method estimated 3 clusters (Supplementary Figure 2), while the silhouette method estimated 2 (SCoef_2_ = 0.23, SCoef_3_ = 0.21). Thus, both 2 and 3 clusters were considered for analysis. Interestingly, forelimb movements of both early blind and sighted animals clustered into similar movement types regardless of whether they were clustered into 2 or 3 stereotypical movements (Supplementary Figure 2). The only observed difference was in the retraction phase of a single movement type of early blind animals (Figure 4E). For both early blind and sighted opossums, the extension phase involved either a quick, medium, or slow placement trajectory.

In whisker-trimmed animals, forelimb movements clustered definitively into two similar types regardless of blind or sighted condition: one movement where the forelimb was lifted off the rung immediately, but took a slow linear trajectory to its peak height, and one quick nonlinear movement with a shallow peak (EB: SCoef_2_ = 0.23, SCoef_3_ = 0.19) (SC: SCoef_2_ = 0.31, SCoef_3_ = 0.17) (Figure 4E). Thus, stereotypical movement types were not influenced by the presence or absence of vision, but instead by the presence or absence of whiskers. Additionally, movements were most notably changed in the retraction phase, showing increased variability and altered waveform shape, illustrating the importance of whiskers in detecting the upcoming rung.

### Opossums adapt to whisker trimming by altering body posture and strategy

Finally, to examine the alternate strategies used by opossums after whisker trimming, we quantified aspects of body posture during correct forelimb placements. First, we quantified the distance in the X direction (stride length) between the right forelimb and right hindlimb, as well as between the right forelimb and snout, during correct motions. Animals with whiskers, regardless of blind or sighted condition, show standard locomotor postures, with limbs oscillating between long (> 5 cm) and short distances (~ 1 cm), reminiscent of the quadrupedal gait cycle (Figure 5A, Figure 5B solid lines). However, whisker-trimmed opossums show flattened trajectories, which never reach large amplitudes (Figure 5A, Figure 5B dotted lines). Instead, these condensed limb distances illustrate the hunched, conservative approach that blind and sighted animals employed when crossing the ladder without whiskers (Supplemental Video 2, Supplemental Video 3).

**Figure 5.**
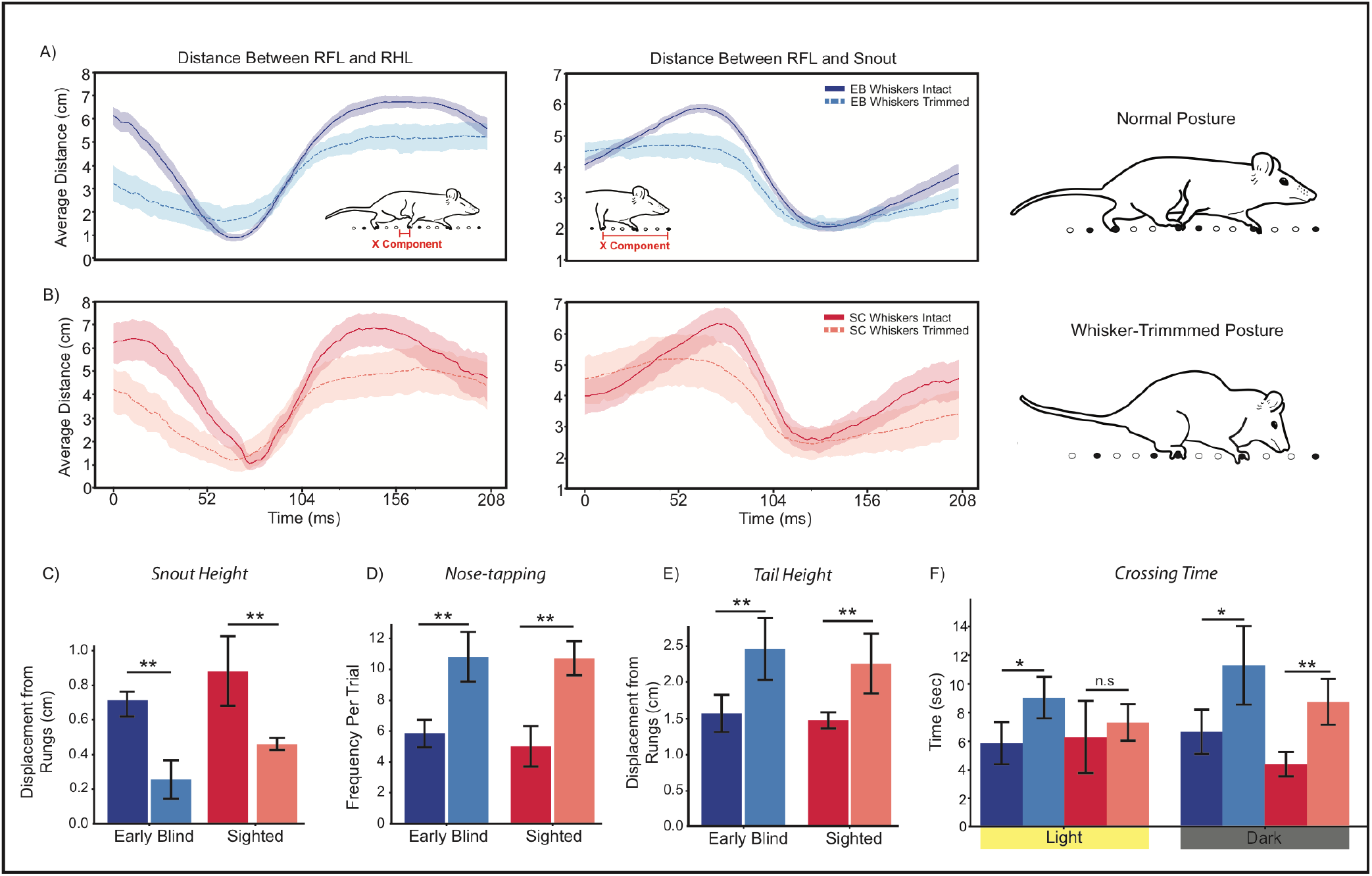
Animals adapt to whisker trimming in similar ways. (A) Line graphs of average stride (x-component only) between the right forelimb and right hindlimb (left) and the right forelimb and snout (right) in early blind (A) and sighted (B) opossums during correct forelimb strikes. In both groups, whisker trimming reduces stride and decreases the distance between the right forelimb and snout. The flattened whisker-trimmed trajectories illustrate the conservative approach taken by whisker-trimmed opossums, as animals hunch their posture and are less willing to take long strides. Illustrations (right) derived from tracings of animals with (top) and without (bottom) whiskers, representative of typical locomotor postures while ladder crossing. (C,D,E) Bar graphs depict quantified aspects of body posture during ladder crossing. (C) Whisker trimming results in significant decreases in average snout height (p =0.004) and (D) more nose tapping behavior (p =0.002), (E) while the tail is held higher (p =0.042). (F) Bar graph shows the interaction between whiskers and lighting condition on crossing time. Whisker trimming results in increased crossing time for early blind animals in the light and dark (p = 0.022, p = 0.015). Whisker trimming only results in increased crossing time for sighted animals in the dark (p = 0.448, p < 0.001).

Additionally, animals without whiskers held their snout closer to the rungs on average (R^2^adj = 0.59, F(4) = 6.46, p = 0.004, ΔEB = −0.46 cm, ΔSC = −0.42 cm) (Figure 5C), and exhibited significantly more nose tapping behavior (R^2^adj = 0.34, F(4) = 6.46, p = 0.002, ΔEB = +3.89 taps, ΔSC = +4.56 taps) (Figure 5D), possibly in order to gain tactile and/or olfactory information. Whisker trimming was also accompanied by an increase in average tail height during correct placements (R^2^adj = 0.51, F(4) = 4.95, p = 0.042, ΔEB = +0.90 cm, ΔSC = +0.78 cm) (Figure 5E), and a dramatic increase in crossing time (R^2^adj = 0.24, F(4) = 7.672, p = 0.001, ΔEB = +4.86s, ΔSC = +3.40s) (Figure 5F).

To further examine how sighted and blind animals adapted to whisker trimming, we analyzed crossing time in both lighting conditions. Proving an effective control, we found that whisker trimming increased crossing time for early blind animals irrespective of lighting condition (p_light_ = 0.022, p_dark_ = 0.015, ΔEB_light_ = +4.00s, ΔEB_dark_ = +5.86s) (Figure 5F). However, whisker trimming only increased crossing time for sighted controls in the dark, suggesting that whisker-trimmed sighted animals employ the use of visual cues when visual information is available (p_light_ = 0.45, p_dark_ < 0.001, ΔSC_light_ = +1.27s, ΔSC_dark_ = +5.44s) (Figure 5F). These metrics indicate that early blind and sighted animals adopt a conservative strategy to ladder crossing when tactile sensory input from the whiskers is removed, and that sighted animals recruit vision in the absence of whiskers.

## Discussion

We quantified the extent to which early vision loss impacts movement strategy and performance on a sensorimotor task involving the spared senses. Our results indicate that early blind (EB) animals have superior performance on the ladder rung task, in part due to increased precision of sensory-guided forelimb placement. Whisker-trimming nullified this advantage, demonstrating that EB animals relied more heavily on input from the whiskers to complete the task than sighted animals (SC). Following whisker trimming, both EB and SC animals adopted similar crossing strategies during the dark condition, but SC animals showed no increase in crossing time in the light condition (while EB animals did), suggesting that sighted animals utilized vision when tactile input from the whiskers was unavailable. We first discuss the increased performance of early blind animals on the ladder rung task and compare this with studies in humans that demonstrate cross-modal behavioral plasticity following the early loss of vision. We then discuss the role of active sensing for locomotion and complex navigational tasks. Finally, we explore the underlying neural mechanisms that may subserve these sensorimotor abilities, and how they have been modified following the early loss of vision.

### Cross-Modal behavioral plasticity following early loss of vision

Preliminary data from our laboratory indicates that EB animals are better at making fine tactile discriminations, but show no differences in whisker angle or whisking frequency compared to SC opossums (Dodson et al., 2017; Ramamurthy & Krubitzer, 2018). In the current study, we found that EB opossums could detect rungs and place their limbs more accurately during ladder rung walking. While few other studies in animals have been conducted, research in humans has extensively examined the behavioral effects of the early loss of vision. Mainly, these studies have focused on two types of behavior: sensory acuity and spatial information processing. For example, it has been shown that EB individuals have increased auditory and tactile spatial acuity, reacting to auditory and tactile spatial targets quicker than SC without changes in performance (for review: Collignon et al., 2009; Collignon & De Volder, 2009). Additionally, Voss and Zatorre found that increases in the thickness of occipital cortex in EB compared to SC subjects correlated with increased performance on pitch and melody discrimination (Voss & Zatorre, 2012). Using a somatosensory spatial discrimination task, Wong and colleagues found that EB individuals have higher tactile acuity on the fingers than SC, but tactile discriminations made with the lips were the same in both groups (Wong et al., 2011). EB individuals also have quicker reaction times and increased accuracy on texture, but not shape discrimination tasks (Gurtubay-Antolin & Rodríguez-Fornells, 2017; Schubert et al., 2017; Wan et al., 2010).

Although blind individuals use input from multiple senses to navigate, most studies of navigation in the early blind focus on the auditory system. Research by Lessard and colleagues found that EB subjects can map the location of auditory tones in space more accurately than SC subjects, and are capable of doing so with information from only one ear (Lessard et al., 1998). More recent studies in humans show that EB echolocators exhibit enhanced auditory spatial navigation abilities compared to EB non-echolocators and sighted controls (Gori et al., 2014; Vercillo et al., 2015). Using a sensory substitution device (SSD) that translated the distance of objects into auditory tones, EB subjects showed equal maze navigation performance to SC subjects once training was complete (Chebat et al., 2015). Another study, which used somatosensory cues relayed through the tongue, found that EB subjects can learn spatial navigation patterns at least equally as well as SC subjects (Kupers et al., 2010). While these studies indicate that cross-modal behavioral plasticity is present in EB individuals, the variations in task parameters and limited research on the strategies that blind individuals use limit our understanding of how early vision loss affects sensory-guided movement using the remaining senses (Klatzky et al., 1995; Passini & Proulx, 1988; Tinti et al., 2006). Given the prevalence of EB individuals who navigate their daily environment with a cane, it is surprising that more research has not been dedicated to SSD’s which focus on tactile input. Moreover, the above studies which do employ tactile information to study spatial information processing in the blind, often used reaction time, and measured spatial acuity based on stimulus detection only. That is, participants in these studies only had to detect a spatial stimulus, but were not required to reach toward or touch the target, neglecting potential differences in movement strategy and motor performance.

An important contribution of the present study is that we not only look at performance, but also quantify the strategy adopted by our different groups when performing the ladder rung task. In a recent review, Schinazi and colleagues stress the importance of studying the relationship between strategy and performance in uncovering differences in spatial navigation abilities between congenitally blind and sighted humans (for review see: Schinazi et al., 2016). In our study, EB opossums had to detect an upcoming rung, accurately place their forelimb on the rung, and maintain a spatial representation of that rung in order to ensure accurate hindlimb placement. We found that EB opossums had greater precision in targeting upcoming rungs with both the fore- and hind limbs while showing minor differences in limb trajectory, thus exhibiting a heightened ability to navigate in a complex tactile space. This superior performance was presumably due to input from the spared senses, such as the whiskers. When this spared sensory input was removed by trimming the whiskers, both EB and SC animals adopted similar crossing strategies, but EB animals showed greater deficits in performance and posture.

Information on the location and distance of gaps is usually provided by the whiskers (Arkley et al., 2017); however, both blind and sighted animals adjusted to whisker trimming by using somatic and possibly olfactory input from the nose, shown by a two-fold increase in nose-tapping behavior. Further quantification of body posture showed that animals in both groups adapted to whisker trimming by taking a more cautious approach - illustrated by a truncated stride length and shortened distance between the snout and forelimb, as well as increased crossing time in most conditions. Of note is that whisker trimming did not cause a significant increase in crossing time for sighted opossums in the light condition, while crossing time did increase for SC in the dark condition, indicating that SC animals may have used the visual system to perform this task when the whiskers were removed. In agreement with Shinazi and colleagues, our results show the importance of studying both performance and strategy, as one or the other may be altered depending on the available sensory information. In our case, without tactile information from the whiskers, EB opossums qualitatively resembled a blind human trying to navigate without a cane, and SC opossums resembled a sighted human trying to navigate in the dark.

### Active sensing is critical for guiding forelimb movements in mammals

Active sensing with the whisker system has been shown to be critical for many animals to detect walls and objects, and is thought to directly guide forelimb placement during locomotion (Arkley et al., 2014, 2017; Grant et al., 2009, 2013; Mitchinson et al., 2007). In the current study, we found that whisker trimming resulted in huge impairments in precision sensory-guided stepping during ladder crossing, regardless of whether opossums were blind or sighted. These results add support to the prevailing theory that whisking is fundamental for guiding limb placement. First and foremost, we found that animals without whiskers had increased forelimb error, but not hindlimb error. Second, in agreement with work on precision stepping in cats and rats (Drew & Marigold, 2015; Whitlock, 2014), where spatial information from the forelimb informs the future trajectory of the hindlimb (by encoding the height and location of obstacles), we found that whisker trimming altered the height of both the forelimb and hindlimb. While both groups of animals exhibited a reduction in forelimb and hindlimb peak height due to whisker trimming, this occurred to a greater extent in EB animals. Moreover, hindlimb trajectory, but not performance, was altered by whisker trimming. Third, recent research in rats has shown that whisker trimming results in increased variation in limb kinematics during locomotion on a continuous substrate (Niederschuh et al., 2015). Similarly, we found that whisker trimming resulted in more variable forelimb movements during the retraction, but not extension phase of the trajectory. We believe the retraction phase to be associated with sensing an upcoming rung, as the nose oscillates vertically during correct placements. While we did not find notable differences in the periodicity of nose oscillations (data not shown), future studies that directly examine the relationship between whisking frequency/contact and forelimb placement will shed light on this aspect of locomotor control. Regardless, increased variance in the retraction but not extension phase may indicate increased hesitation when opossums are detecting an upcoming rung’s position, but once located, can make a precise placement.

Across metrics, we found that EB animals show a trend for lower variation. While this speaks to the increased precision of whisker-guided forelimb movements in blind opossums, we cannot rule out visual input altering attention in sighted animals. Nevertheless, in the presence or absence of vision, forelimb movement types cluster into similar stereotyped movements. On the other hand, we found that the absence of whiskers forces forelimb placement trajectories to conform to two similarly stereotyped movements in blind and sighted opossums. Thus, it was the loss of whiskers, and not the loss of vision, that impacted the produced stereotyped forelimb movements. This highlights the remarkable flexibility in behavior when sensory input is altered during development, and the compensatory nature of this process, where heightened performance on some tasks is accomplished via increased reliance on spared sensory systems.

### Neural mechanisms that may subserve adaptive cross modal behavioral plasticity

Following the early loss of vision, cortical areas associated with both the lost sense as well as spared sensory systems are profoundly affected. In EB mice, rats, and cats, the connections of primary visual cortex are drastically altered, such that cortico-cortical projections from somatosensory and auditory areas densely project to what would have been visual cortex (Berman, 1991; Dye et al., 2012; Laemle et al., 2006; Négyessy et al., 2000). It should be noted that the developmental time when vision is lost strongly impacts the extent of anatomical changes that occur (Mezzera & López-Bendito, 2016), with the greatest effect occurring prior to the onset of thalamo-cortical afferentation. Studies in our laboratory demonstrate that when opossums are enucleated prior to the onset of spontaneous activity from the retina and before the establishment of retino-geniculo-cortical connections (P4), visual cortex receives input from somatosensory and auditory structures of the cortex and thalamus (Karlen et al., 2006). The early loss of vision also results in increased connectivity within somatosensory cortex, as well as between somatosensory and visual cortex, and somatosensory and posterior parietal cortex (Dooley & Krubitzer, 2019; Kozanian et al., 2015). Similar to studies in animal models, indirect measures of connectivity, such as resting state fMRI and DTI, in EB humans shows that S1 is more densely connected (correlated) with V1, and that posterior parietal cortex exhibits denser projections with sensory areas as well (Klinge et al., 2010; Liu et al., 2007; Ptito et al., 2005, 2008; Shu et al., 2009; Wittenberg et al., 2004).

Functional changes accompany these anatomical changes in early blind animals and humans. Visually deprived mice and rats have altered excitatory synaptic function in V1, and neurons in a large proportion of the reorganized primary visual cortex respond to auditory stimuli (Goel et al., 2006; Piché et al., 2007; Zheng et al., 2014). In early blind *Monodelphis*, all of what would be visual cortex (V1) is co-opted by the somatosensory and auditory system (Kahn & Krubitzer, 2002). Importantly, receptive fields from neurons in the re-organized V1 are concentrated on the vibrissae and face. There are only a few studies that examined the functional organization of somatosensory cortex following early loss of vision. Using single-unit electrophysiology, Toldi and colleagues found that receptive fields in the barrel cortex of rats were significantly altered by early vision loss (József Toldi et al., 1994). Another study in mice used cytochrome oxidase staining as an indirect measure of functional organization, and found that early vision loss resulted in an increase in the size of barrel cortex (Rauschecker et al., 1992). Recent work from our laboratory found that neurons in the whisker-representation of S1 in EB opossums exhibit spatial sharpening of receptive fields and improved population decoding for whisker-stimulus position (Ramamurthy & Krubitzer, 2018), indicating that individual neurons in S1 in EB opossums have better discriminability. While the functional organization of posterior areas in parietal cortex has not been extensively explored in *Monodelphis*, research in rats and cats shows that activity in posterior parietal cortex increases before and during gait modifications (Beloozerova & Sirota, 2003; Whitlock, 2014). Importantly, in blind rats, neurotoxic lesions to posterior parietal cortex cause deficits in spatial memory during maze-running (Pinto-Hamuy et al., 2004).

Functional changes due to early vision loss have been extensively studied in humans. In EB individuals, V1 is activated during somato-motor tasks involving the hands (Gizewski et al., 2003), and during tasks involving memory, spatial processing, and language (Amedi et al., 2003; Bedny et al., 2011; Ricciardi et al., 2014). Visual cortex is also activated by guided hand motions, regardless of if participants are SC or EB (Fiehler & Rosler, 2010). However, during this task, EB individuals also showed activation of auditory and extrastriate cortex, when SC subjects did not. Further, in congenitally blind humans, posterior parietal cortex retains its role as an encoder of the spatial position of a reach target and shows increased integration of tactile information (through variability of the BOLD signal) in participants completing a tactile spatial discrimination task (Leo et al., 2012; Lingnau et al., 2014).

Taken together, studies in animal models and humans demonstrate that visual cortex is not dysfunctional in the absence of vision, but instead contributes to a number of behaviors including tactile and spatial processing (for review: Ricciardi et al., 2014). These studies also suggest that superior performance on tactile discriminations may be due to changes in the structure and function of somatosensory and posterior parietal cortices. Given previous data from our laboratory in EB opossums showing: 1) a somatosensory-driven reorganization of V1, 2) projections from somatosensory cortex to V1, 3) alterations in neural response properties in the whisker representation in S1, and 4) increased connections between S1 and posterior parietal cortex, we posit that these neural changes could support the heightened abilities of EB opossums observed in this study, and also explain the extreme detriments to performance when the whiskers were trimmed. While we have focused on the cortical mechanisms that may account for differences in performance between EB and SC animals, it is possible that the morphology of the whiskers may also be altered, as seen in visually deprived mice and cats, who have increased diameters of macro-vibrissae (Rauschecker et al., 1992). Importantly, these data have implications for congenitally blind humans, suggesting that sensory substitution devices which rely on tactile input, and behavioral therapies which involve the rapid acquisition of spatial information through tactile input, will be effective in increasing the ability of EB individuals to navigate in a complex environment.

## Materials and Methods

### Subjects

Twenty-seven adult (≥ 180 days) short-tailed opossums (*Monodelphis domestica*) were used to test variable ladder rung walking. Twelve animals (5 male, 7 female) were bilaterally enucleated on postnatal day 4 (P4), while fifteen animals (8 male, 7 female) were sighted littermate controls. Four animals from each group were also used for limb tracking analysis. All animals were obtained through our breeding colony at the University of California, Davis. Animals were reared in standard laboratory conditions, weaned at 2 months, and separated from littermates at 4 months. Upon separation, females were co-housed with one female littermate, while males were single-housed. All experimental procedures were approved by UC Davis IACUC and conform to NIH guidelines.

### Bilateral Enucleation Surgery

Bilateral enucleations on *Monodelphis* pups in our laboratory have been described in detail previously (Kahn & Krubitzer, 2002; Karlen & Krubitzer, 2009). In brief, on postnatal day 4 (P4), mothers were first anesthetized with isoflurane (5%), and anesthesia was maintained with Alfaxalone (3 mg/kg, 10 mg/ml IM). Pups were anesthetized via hypothermia. At this age, pups are fused to the mother’s nipple, and do not detach until around the third postnatal week. Under microscope guidance, the skin covering the retina was incised and retracted and the immature retina was removed. A flush of sterile saline was then applied to rinse the area, and the skin was replaced and sealed with surgical glue. Approximately half of each litter was bilaterally enucleated in this manner, while the other half served as littermate controls. Once enucleation procedures were complete, mothers were allowed to recover in their home cage.

### Behavioral Testing

When enucleated and control animals reached maturity (≥ 180 days), we tested their performance on the variable ladder rung walking task, commonly used in rodents (Schönfeld et al., 2017). Animals were first tested in both light and dark conditions (in varying order) with their whiskers intact. We used a 660nm lamp (outside the peak-sensitivity range for cones in *Monodelphis*) to ensure accurate video collection and scoring in the dark condition (Hunt et al., 2009; Seelke et al., 2014). At least one week after initial testing, all mystacial and genal whiskers were trimmed and animals were tested again in both lighting conditions on two consecutive days. To trim the whiskers, animals were briefly anesthetized with isoflurane (2-5%) and macrovibrissae were trimmed down to ~1 mm using an electric clipper. Animals were allowed to recover for 12 hours in their home cage before undergoing testing. All whisker-trim testing was conducted within 72 hours of trimming to prevent animals from recovering whisker sensation due to significant regrowth.

### Variable Ladder Rung Walking

The ladder rung apparatus consisted of an elevated trough with a floor of ladder rung-like pegs on which the animals could walk (Figure 1A). The trough was constructed from two Plexiglas walls 1 meter in length, connected by a floor of metal pegs (3 mm diameter, 10 cm wide) similar to that used by Metz and Whishaw (Metz & Whishaw, 2009). Holes for the removable rungs forming the floor of the apparatus were spaced at 1cm intervals. The apparatus was supported by a neutral start box on one end and the animal’s home cage on the other end, and was raised approximately 1 meter from the floor to discourage jumping. Animals crossed the apparatus from the neutral start cage to their home cage. Motivation to reach the home cage was high, so no additional reward was given.

For the baseline (standard pattern) condition, all rungs were equally spaced 2 cm apart. For variable patterns, rungs were spaced 1 - 5 cm apart, with no more than 3 rungs of 1 cm spacing in a row before a 2 cm gap. Variable patterns were generated using a custom Python program which determined random 10-rung patterns, which were then repeated over the 1-meter long apparatus. To capture naturalistic behavior, without effects of training, each animal participated in only five days of testing. Day 1 consisted of five runs of the standard pattern to allow for habituation, with all rungs at 2 cm spacing. Days 2 - 5 consisted of three standard patterns per animal, followed by two or three variable patterns. Two consecutive days of testing were done with lights on (either days 2&3 or days 4&5) on variable patterns (light condition), and two consecutive days of testing were done in 660 nm red light (dark condition). Trials were video-recorded and scored (see below). Trials in which animals did not cross at least half (0.5 m) of the apparatus before stopping or reversing direction were not scored (< 5% of total trials).

### Scoring

We adapted the scoring method originally used by Metz and Whishaw (Metz & Whishaw, 2009). The four-point scale included: (0) Total Miss: 0 points were awarded when a limb entirely missed a rung, and body posture was disturbed. (1) Slip: 1 point was awarded when the limb initially contacted a rung before slipping off. (2) Correction/Replacement: 2 points were awarded when the limb aimed for one rung, but was placed on a different rung before contacting the first, or when the limb was placed on one rung, but moved to another rung before weight-bearing. (3) Correct Placement: 3 points were awarded when a limb was advanced and placed on a rung, and could bear weight without causing a visible disturbance in body posture. Scores of (0) and (1) were combined as errors, and divided by the total number of attempts to produce an error rate (Figure 1A). Crossing time was also scored, excluding time when the animal would stop to groom.

Because our initial observations showed animals changing head pitch to tap their nose on the next rung before stepping, we quantified the frequency of nose taps in each trial as state events, when the rostrum made physical contact with a rung. Scores for foot fault, crossing time, and nose taps were averaged by animal across all variable patterns.

### Manual video analysis

Videos were recorded with various cameras to allow for wide-field, standard, and high-speed video capture from multiple angles. A Canon VIXIA HF R500 camcorder (60 fps) and GoPro Hero 6 (240 fps) were used to record light trials, while a SEREE FHD 1080P camcorder (30 fps), was used to record dark trials. Videos were scored in VLC Player using frame-by-motion, and at 25 and 50% of standard playback speed (7.5 – 30 fps) by independent raters and/or using the DeepLabCut analysis system (see below). Two observers blind to the animal’s condition scored all trials independently. Scores were averaged between observers to provide a single measure (for inter-rater reliability measures see Statistics; Supplementary Figure 1).

### Automated video analysis

Video of ladder rung crossing strategy was also analyzed by DeepLabCut, a marker-less pose-detecting machine learning Python package (Mathis et al., 2018). This toolbox utilizes Google Tensorflow and ResNet to track any user-defined body part, animal, or object in successive video frames. DeepLabCut is blind to experimental condition, and is capable of tracking movements that allow us to quantify and analyze differences in strategy. We trained the neural network for 400,000 iterations on 300 labeled frames. This regimen was adequate to produce the desired fit of the model to the training data (loss <.005), such that predictions of movements by DeepLabCut resulted in accurate tracking of body parts (Supplemental Video 1). Custom Python programs were produced to analyze per-frame positional information of all four limbs, the tail, and snout, during light and dark trials (code available at: https://www.github.com/maceng4/Monodelphis_Ladder_Rung).

To study changes in locomotor patterns as opossums crossed the ladder apparatus, we restricted our analysis to correct placements of the right forelimb, during which: 1) no other limbs were currently making an error and 2) all DLC trackers accurately labeled the assigned body part for the duration of the gait cycle. Both a human observer and computer vision (continuous plots of each tracker) were used to provide confirmation. Correct strikes were defined as motions of the right forelimb which began on a single rung and involved reaching for and correctly grasping a new rung. These motions corresponded to performance scores of 3: correct placement. We selected 100 video frames centered on the peak of each forelimb movement using the above criteria (50 frames prior to the peak and 50 frames after, captured at 120 × 4 fps). Consequently, each analyzed strike yielded 208 milliseconds of data of the whole body during a correct placement of the right forelimb.

Data across all strikes was aligned to the highest peak during forelimb motion, aggregated along the time axis (208 milliseconds) and separated into × and Y displacement components to provide average trajectories per group. The Y-component represents height (displacement from the ladder apparatus in centimeters) and the X-component represents step length (Figure 1B). To determine differences between groups, we characterized trajectories by quantifying peak width at half-height, average peak height, average displacement, and binned variance. We accounted for differences in animal size by scaling all data for an individual animal by its average forelimb height during a correct movement, before combining data across animals. Statistical models tested for differences between experimental conditions using per-animal averages.

For analyses of stereotyped movements (K-means clustering), we generated scripts which implemented supervised machine learning packages from sci-kit learn (sklearn). We used the elbow method in concert with the silhouette coefficient (described below) to select the number of clusters (movement types) that a given forelimb movement could be assigned to (Supplemental Figure 2) (Syakur et al., 2018; Zhou & Gao, 2014). For the elbow method, the x-value at which exponential decay ceases (i.e at the elbow joint of the line graph) estimated the optimal number of clusters to use for a given dataset. To confirm this estimation, we then used the silhouette coefficient, calculated using the mean intra-cluster distance and the mean nearest-cluster distance. In short, the silhouette coefficient is a measure of separation between groups, where a high number indicates an instance is well-matched to its own cluster. The two highest silhouette coefficients were used to select the number of clusters per experimental condition. In our case, this resulted in either 2 or 3 clusters for each condition (see Results; Supplementary Figure 2).

### Statistical analysis

Statistical tests were performed using custom Python scripts (https://www.github.com/maceng4/Monodelphis_Ladder_Rung). We generated fixed effects linear models and used analyses of variance to assess differences between groups. For all analyses, we tested multiple models using backward selection to ensure statistical accuracy. To assess differences in performance between early blind and sighted animals, we used a fixed effects linear model, testing for main effects of sightedness, lighting, biological sex, and the presence of whiskers. This model included interaction terms for the presence of whiskers (trimmed or intact) and lighting condition (light or dark), while sex was included as a covariate (see below: full model). After finding no significant main effect for lighting condition on performance (Supplemental Figure 1C), we collapsed across light and dark conditions for further analysis (see reduced model).

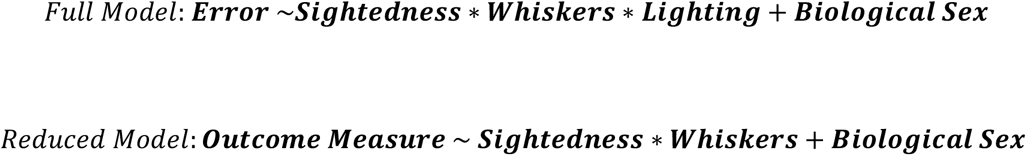

We used repeated measures ANOVAs to test for differences within and across testing days, and chose an alpha level < .05 for significance, and the Holm-Sidak method to correct for multiple comparisons. Bar graphs and trajectory data are presented as means with bootstrapped 95% confidence intervals. Given the variability we observe when working with wild-type strains of laboratory animals on untrained naturalistic behavioral tasks (Englund et al., 2018), we tested for normalcy in performance data using the Shapiro-Wilk test. Indeed, scores for total error were found to be normally distributed (n = 27, w = 0.979, p = 0.13), as was trajectory data of forelimb peak height gathered from the subset of animals used for DLC tracking (n = 8, w = 0.959, p = 0.648).

To test for inter-rater reliability we first generated a linear model, finding significant agreement between raters (R^2^adj = 0.712, F(1) = 174.2, p < 0.001; Supplemental Figure 1B). Additionally, we calculated the intraclass correlation, also finding good agreement (Single fixed raters: ICC3 = 0.84, p < 0.001). This test is commonly used to assess consistency when quantitative measurements (in our case error percentage) are made by different observers (Hallgren, 2012).

## ACKNOWLEDGEMENTS

Thanks to Dr. Andrew Fox and Dr. Eliza Bliss-Moreau for providing consultation on statistical measures and python scripts. Thanks to Cynthia Weller, Heather Dodson, and Carly Jones for assistance with data collection; and to Dr. Andrew Halley, Dr. Deepa Ramamurthy, Dr. Chris Bresee, and Carlos Pineda for manuscript comments. This research was supported by the McDonnell Foundation (Grant 220020516 to L.A.K).

## SUPPLEMENTAL MATERIAL

**Supplemental Figure 1.**
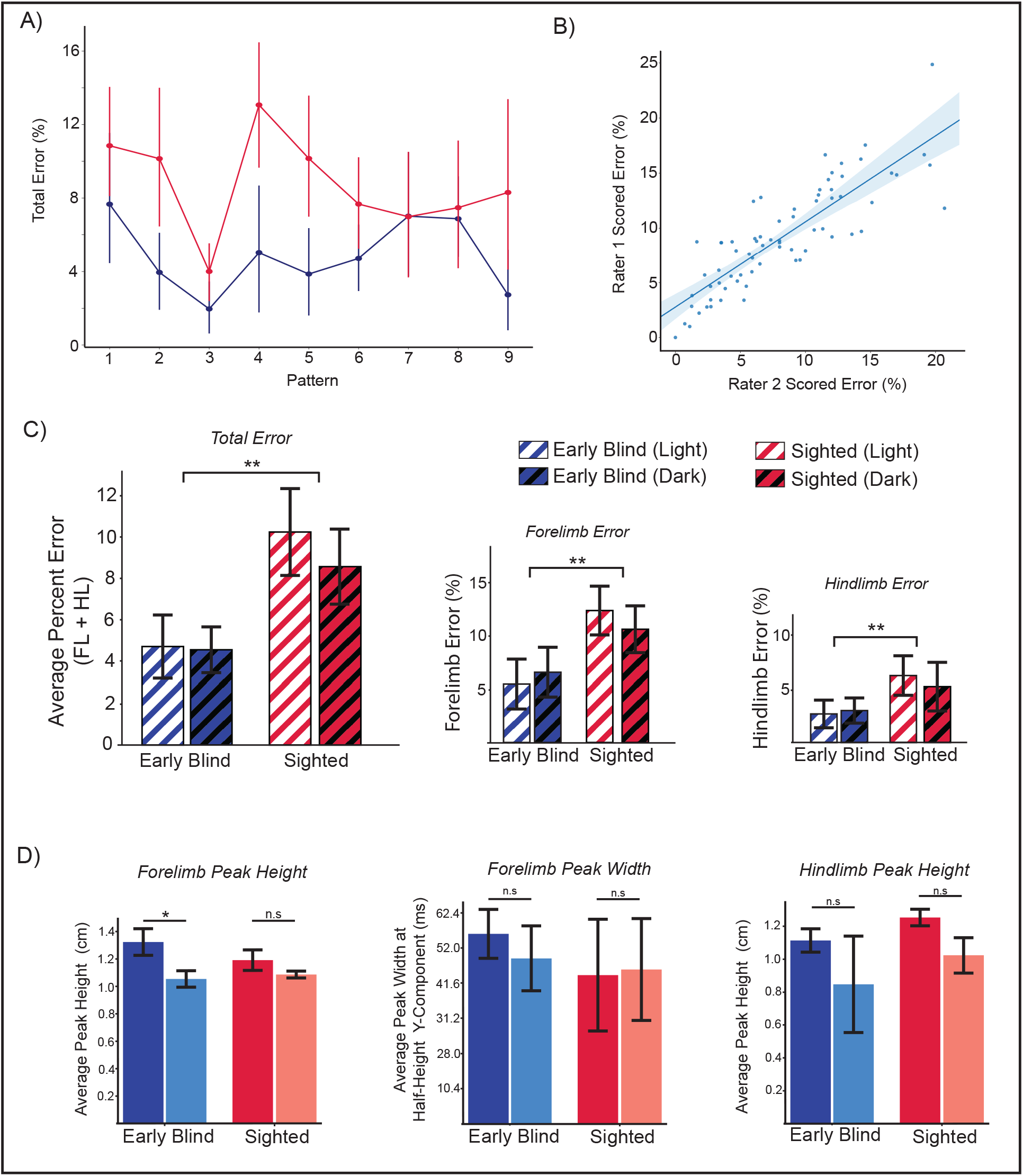
(A) Average error by rung placement pattern in early blind (blue) and sighted (red) opossums. Whisker-trimmed conditions are represented by lighter shades. Early blind animals exhibit less error on average on 8 out of 9 patterns. We generated a linear model testing for the effects of pattern and order of pattern completion (dark first or light first) on total error, and found that neither significantly predicted rung walking error (R2adj = .086, F(3) = 7.997, p = .367, p = .207). (B) Scatterplot with fit line showing the degree of agreement between Rater 1 and Rater 2 on scoring variable ladder rung walking (R2adj = .712, F(1) = 174.2, p < .001). Interclass correlation was used to measure inter-rater reliability (Single fixed raters: ICC3 = 0.84, p < .001). (C) Average rung walking error in light (white stripes) and dark (black stripes) lighting conditions. Under the full model (see methods) no significant main effect was found for lighting condition (R2adj = .32, F(8) = 6.624, p = .718). (D) Bar graphs depicting quantification of limb trajectories. (Left) Average forelimb peak height is significantly reduced in early blind (R2adj = .701, F(2) = 8.03, p = .016) but not sighted opossums (R2adj = .236, F(2) = 2.24, p = .119) after whisker trimming. (Middle) Forelimb peak width at half height is not significantly altered due to whisker trimming (R2adj = −.307, F(4) = 0.973, p = .696). (Right) Hindlimb peak height is not significantly altered due to whisker trimming (R2adj = .234, F(4) = 2.144, p = 0.083).

**Supplemental Figure 2.**
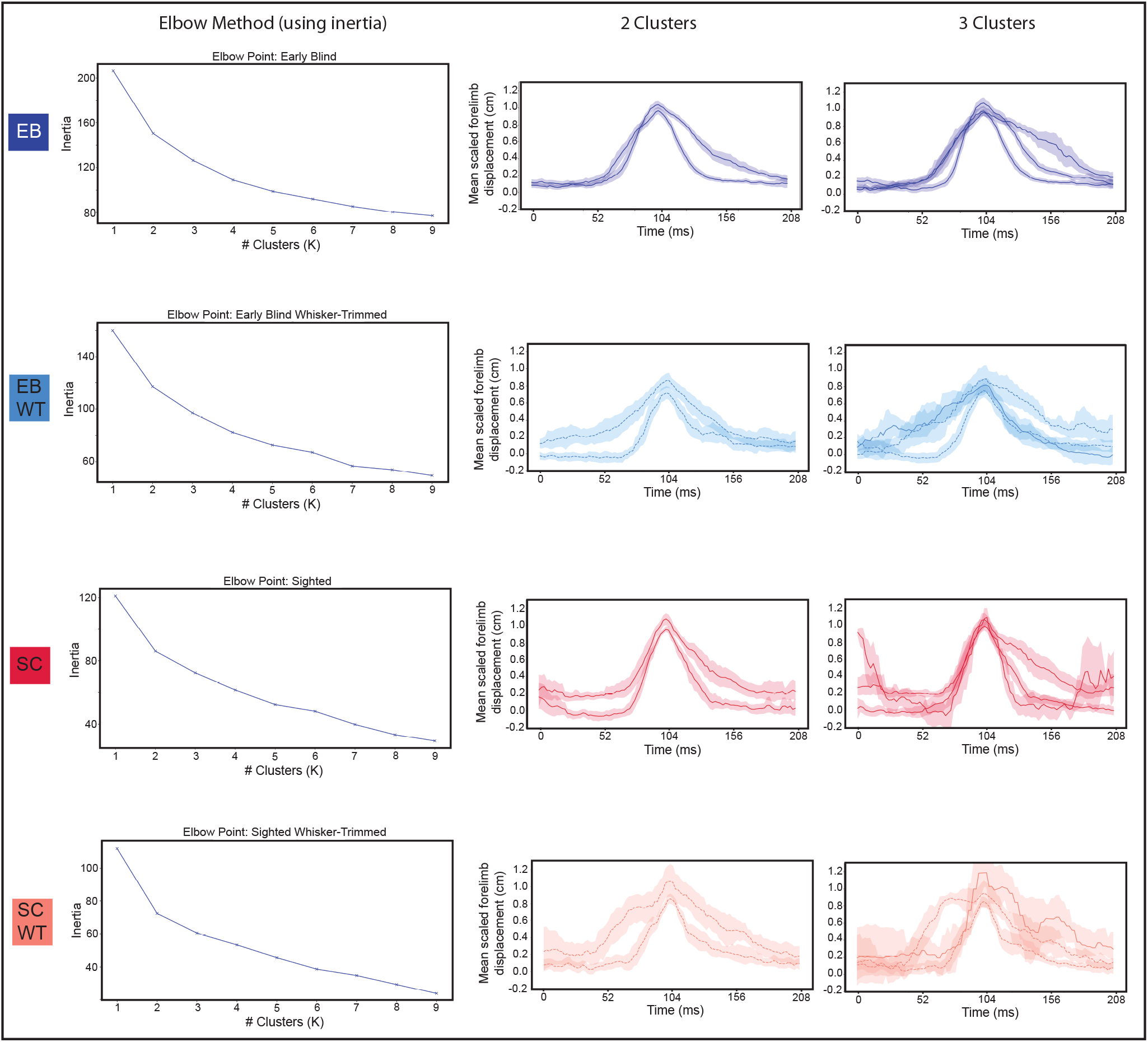
K-Means clustering provides stereotypical forelimb movement types during correct placements. Column 1: The elbow point method and silhouette coefficient were used to determine the number of clusters, and thus the number of stereotyped movements for a given experimental condition. We chose the number of clusters after exponential decay ceased, and/or the silhouette coefficient was closest to 1. Columns 2 and 3: All movements were assigned to both 2 and 3 clusters to illustrate how movements change before and after whisker trimming, regardless of cluster number. Column 2: When 2 clusters are used, the most notable difference is seen in the first half of one movement type (0 – 104ms) in whisker-trimmed conditions, where the forelimb is raised, but takes a linear trajectory to the peak. The other type of movement is noted to be a shallow and quick step. Column 3: When 3 clusters are used to stereotype movements, this difference is still present.

